# Distinct SARS-CoV-2 Antibody Responses Elicited by Natural Infection and mRNA Vaccination

**DOI:** 10.1101/2021.04.15.440089

**Authors:** Rafael Assis, Aarti Jain, Rie Nakajima, Al Jasinskas, Saahir Kahn, Anton Palma, Daniel M. Parker, Anthony Chau, Amanda Leung, Christina Grabar, Fjolla Muqolli, Ghali Khalil, Jessica Colin Escobar, Jenny Ventura, D. Huw Davies, Bruce Albala, Bernadette Boden-Albala, Sebastian Schubl, Philip L. Felgner

## Abstract

We analyzed data from two ongoing COVID-19 longitudinal serological surveys in Orange County, CA., between April 2020 and March 2021. A total of 8,476 finger stick blood specimens were collected before and after an aggressive mRNA vaccination campaign. IgG levels were determined using a multiplex antigen microarray containing 10 SARS-CoV-2 antigens, 4 SARS, 3 MERS, 12 Common CoV, and 8 Influenza antigens. Twenty-six percent of 3,347 specimens from unvaccinated Orange County residents in December 2020 were SARS-CoV-2 seropositive. The Ab response was predominantly against nucleocapsid (NP), full length spike and the spike S2 domain. Anti-receptor binding domain (RBD) reactivity was low and there was no cross-reactivity against SARS S1 or SARS RBD. An aggressive mRNA vaccination campaign at the UCI Medical Center started on December 16, 2020 and 6,724 healthcare workers were vaccinated within 3 weeks. Seroprevalence increased from 13% in December to 79% in January, 93% in February and 99% in March. mRNA vaccination induced much higher Ab levels especially against the RBD domain and significant cross-reactivity against SARS RBD and S1 was also observed. Nucleocapsid protein Abs can be used to distinguish individuals in a population of vaccinees to classify those who have been previously infected and those who have not, because nucleocapsid is not in the vaccine. Previously infected individuals developed higher Ab titers to the vaccine than those who have not been previously exposed. These results indicate that mRNA vaccination rapidly induces a much stronger and broader Ab response than SARS-CoV-2 infection.

## Introduction

Protective efficacy of SARS-CoV-2 spike mRNA vaccines reported by the developers, Pfizer and Moderna, has been successful, showing convincing evidence of protection as short as 14 days after the first immunization [1, 2]. This timeframe is similar to the observed seroconversion times of natural infection that ranges from 10 to 14 days [3, 4]. However, in contrast to the vaccine, it is not yet clear how protective the antibodies induced by natural infection are and how long the protection will last as reports have shown that antibodies generated in response to the infection wane after a few months and can reach baseline levels before the first year [4].

To further understand the mRNA vaccine induced immune response we were interested to compare the antibody response induced by the vaccine with that induced by natural exposure to SARS-CoV-2. Here we show results using a multiplex solid phase immunofluorescent assay for quantification of human antibodies against 37 antigens from SARS-CoV-2, other novel and common coronaviruses, and influenza viruses that are causes of respiratory infections (Figure 1) [5–9]. This coronavirus antigen microarray (COVAM) assay uses a small volume of blood derived from a finger stick, does not require the handling of infectious virus, quantifies the level of different antibody types in serum and plasma and is amenable to scaling-up. Finger stick blood collection enables large scale epidemiological studies to define the risk of exposure to SARS-CoV-2 in different settings.[10] Since the assay requires 1 microliter of blood it is also practical for monitoring immunogenicity in neonates, children and small animal models.

**Figure 1.**
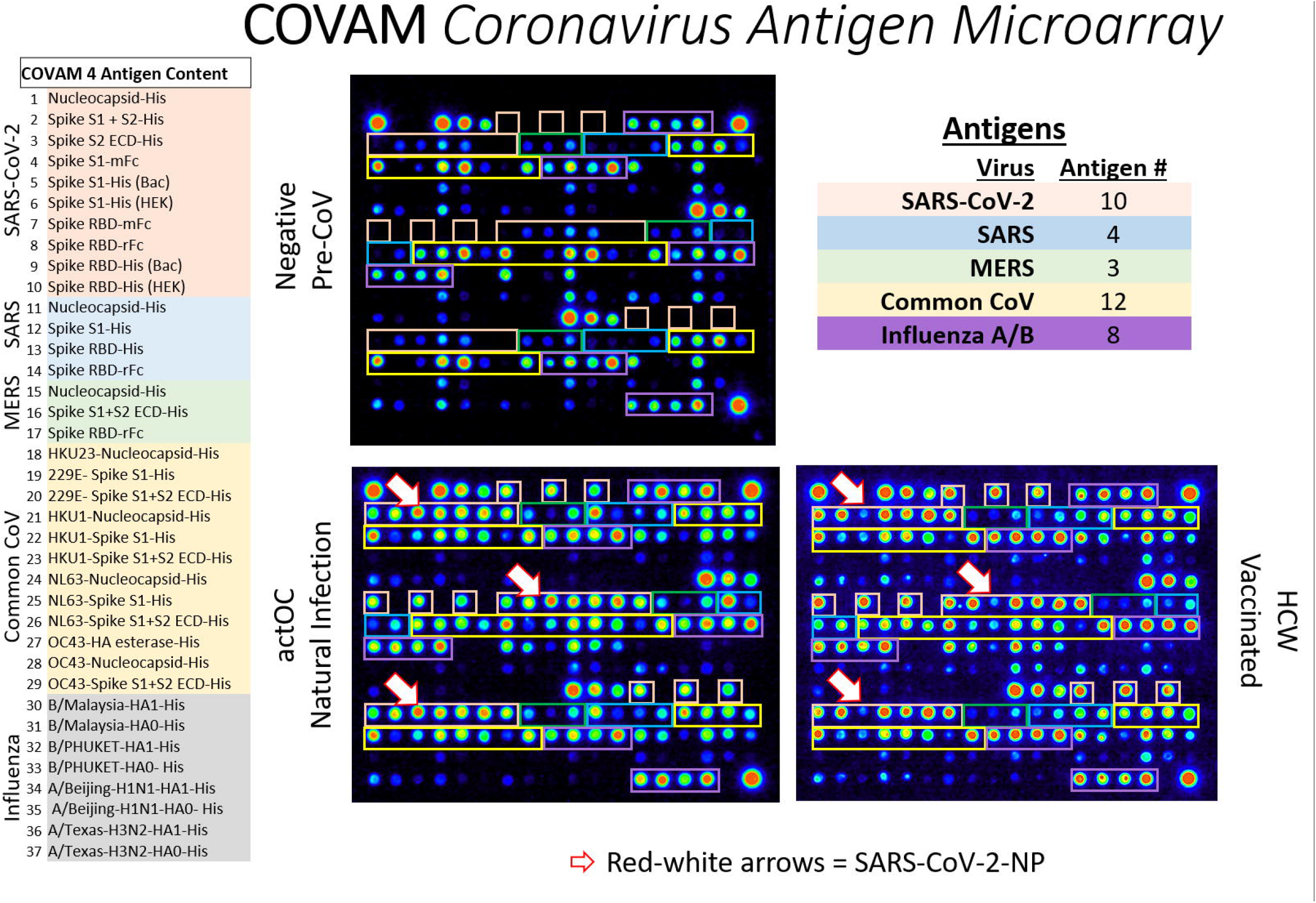
The content of the Coronavirus Antigen Microarray is shown. There are 10 SARS-CoV-2 antigens, 3 SARS, 3 MERS, 12 Common COV, and 8 influenza antigens. Each antigen is printed in triplicate and organized as shown on the images with Orange boxes around the SARS-CoV-2 antigens, Blue SARS, Green MERS, Yellow Common CoV, and Purple for Influenza. Three different samples are shown, a Negative Pre-CoV, Natural Infection (actOC), and a sample from an mRNA vaccinee (HCW). The Pre-CoV sample has negligible reactivities to SARS-CoV-2, SARS and MERS, whereas Natural Infection and the vaccinees have significant Abs against the novel CoV. The red-white arrows point to the nucleocapsid protein which detects antibodies in naturally exposed people but not in the vaccinees.

Our results show that mRNA vaccines are remarkably effective at elevating Ab levels against SARS-CoV-2 antigens, rapidly converting seronegative individuals to seropositive. The observed seroconversion level and breadth across diverse coronavirus strains induced by the mRNA vaccines is much greater than that induced by natural infection. After probing more than 8,729 pre- and post-vaccination specimens our results confirm that the mRNA vaccines can be used in an aggressive and targeted vaccination campaign to immunize large groups within a matter of weeks.

## Results

### mRNA vaccination achieves 99% seropositivity within 3 months after initiating an aggressive and inclusive vaccination campaign

This study was designed to track the seroprevalence at UCIMC since May 2020 and in the Orange County community that is served by the hospital system starting in July (Table 1). In July the observed seroprevalence in Santa Ana zip codes was 18%, and in December it increased to 26% (Figure 2A). Prior to the vaccination campaign in December 2020, the seroprevalence at the UCIMC reached 13%, half of the prevalence measured in Santa Ana. This observation suggests that strict transmission control measures enforced at the hospital played a role in keeping COVID-19 exposure levels low. On December 16, 2020 the vaccination campaign started at the hospital and seroprevalence for the UCIMC population jumped from 13% (early December) to 78% in January, 93% in February, and 98.7% in the last week of March 2021 (Figure 2B). This observation strongly corroborates the high efficacy of the nucleic acid vaccine in stimulating an antibody response and also highlights the success of the vaccination campaign that immunized 6724 HCW from 12/16/2020-1/05/2021, and 10,000 more since then.

**Table 1:** Study Design. Finger stick blood specimens were collected at weekly intervals from drive-through locations around Orange County and from healthcare workers at the University of California Medical Center. Individual samples were probed on the COVAM, quantified and analyzed. Personalized serology reports were generated and linked to individual QR codes for everyone to access their own report.

Samples tested
| <u>Collection</u> | <u>Number</u> | <u>Date</u> |
| --- | --- | --- |
| Orange County | 2,979 | July '20 |
| Santa Ana | 3,347 | Dec '20 |
| UCI Healthcare Workers | 1,060 | May '20 |
| UCI Healthcare Workers | 313 | Dec '20 |
|  |  |  |
| <u>Vaccination Start Date</u> | <u>December 16, 2020</u> |  |
| UCI Healthcare Workers | 140 | Jan '21 |
| UCI Healthcare Workers | 750 | Feb '21 |
| UCI Healthcare Workers | 157 | Mar '21 |
| Total | 8,746 |  |

Measurements
| <u>Virus</u> | <u>Antigen #</u> |
| --- | --- |
| SARS-CoV-2 | 10 |
| SARS | 4 |
| MERS | 3 |
| Common CoV | 12 |
| Influenza A/B | 8 |
| Total | 37 |
| Triplicate | 111 |
| IgG&IgM | 222 |
|  |  |
| Specimens | 8,746 |
| Measurement# | 1,941,612 |

**Figure 2.**
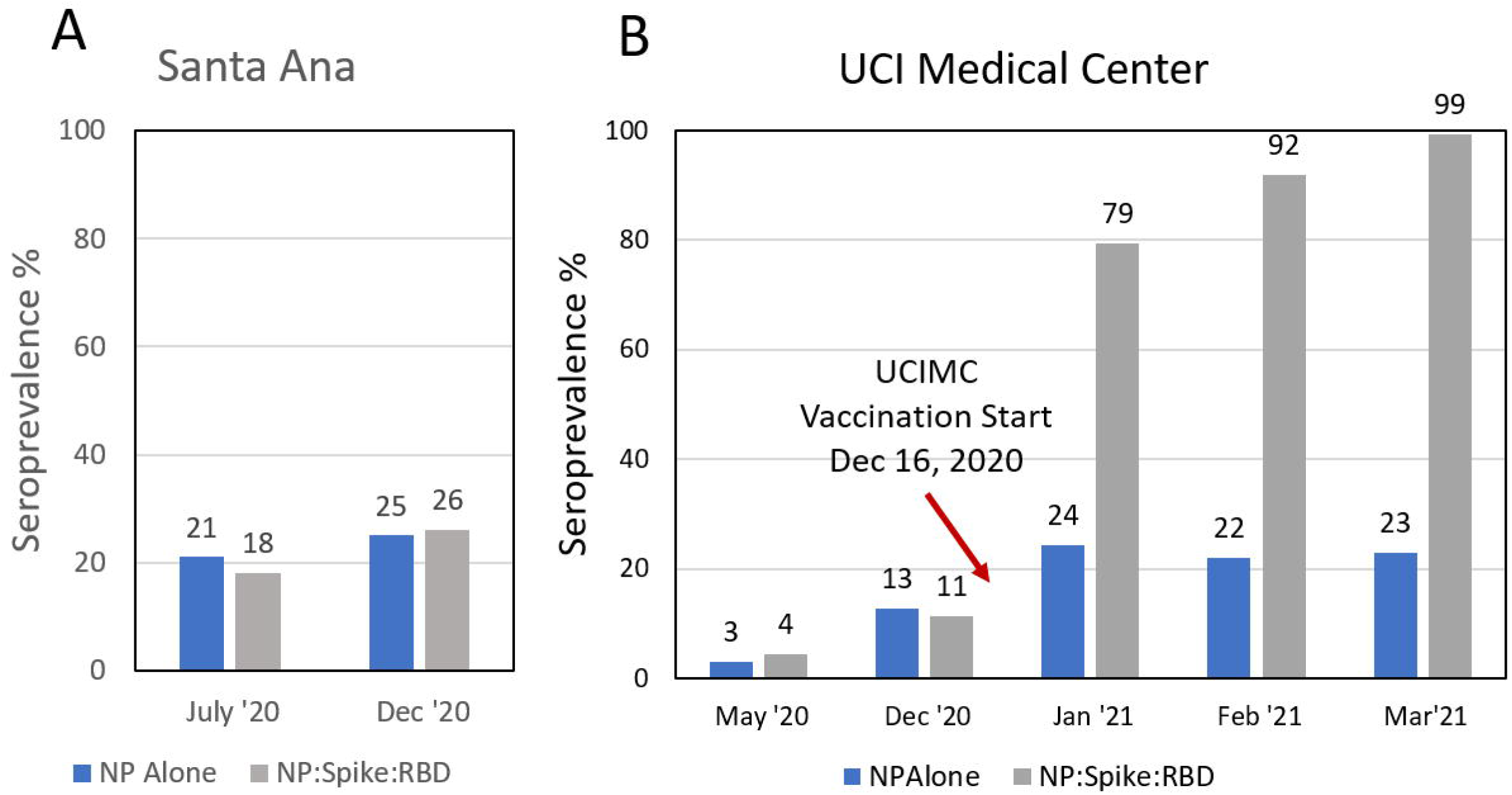
**A**. Finger stick blood specimens were collected from Orange County in July (2,979 specimens) and Santa Ana in December (3,347 specimens), and seroprevalence measured on the COVAM array. **B**. Seroprevalence in cross-sections from the UCI Medical Center was measured by COVAM analysis in May and December before the start of the mRNA vaccination campaign on December 16, 2020 and monthly post vaccination time points in 2021. The gray bar is the COVAM seroprevalence prediction and the blue bar is the nucleocapsid protein seropositivity.

In contrast, comparing the reactivity to the SARS-CoV-2 antigens, differences were noted in the Ab responses induced by the vaccine compared to natural exposure. (Figure 2). The nucleocapsid protein is an immunodominant antigen for which the antibody response increases in concordance with natural exposure (Figure 2A, 3A and 4). However, nucleocapsid is not a component of the mRNA vaccines and consequently there is no vaccine-induced increase in Ab against this antigen. Accordingly, anti-spike antibody levels increased in vaccinees while the nucleocapsid protein Ab level remained constant between Jan and March 2021. (Figure 2B) This suggested that anti-nucleocapsid antibodies can be used as a biomarker of prior natural exposure within a population of seropositive vaccinees.

**Figure 3.**
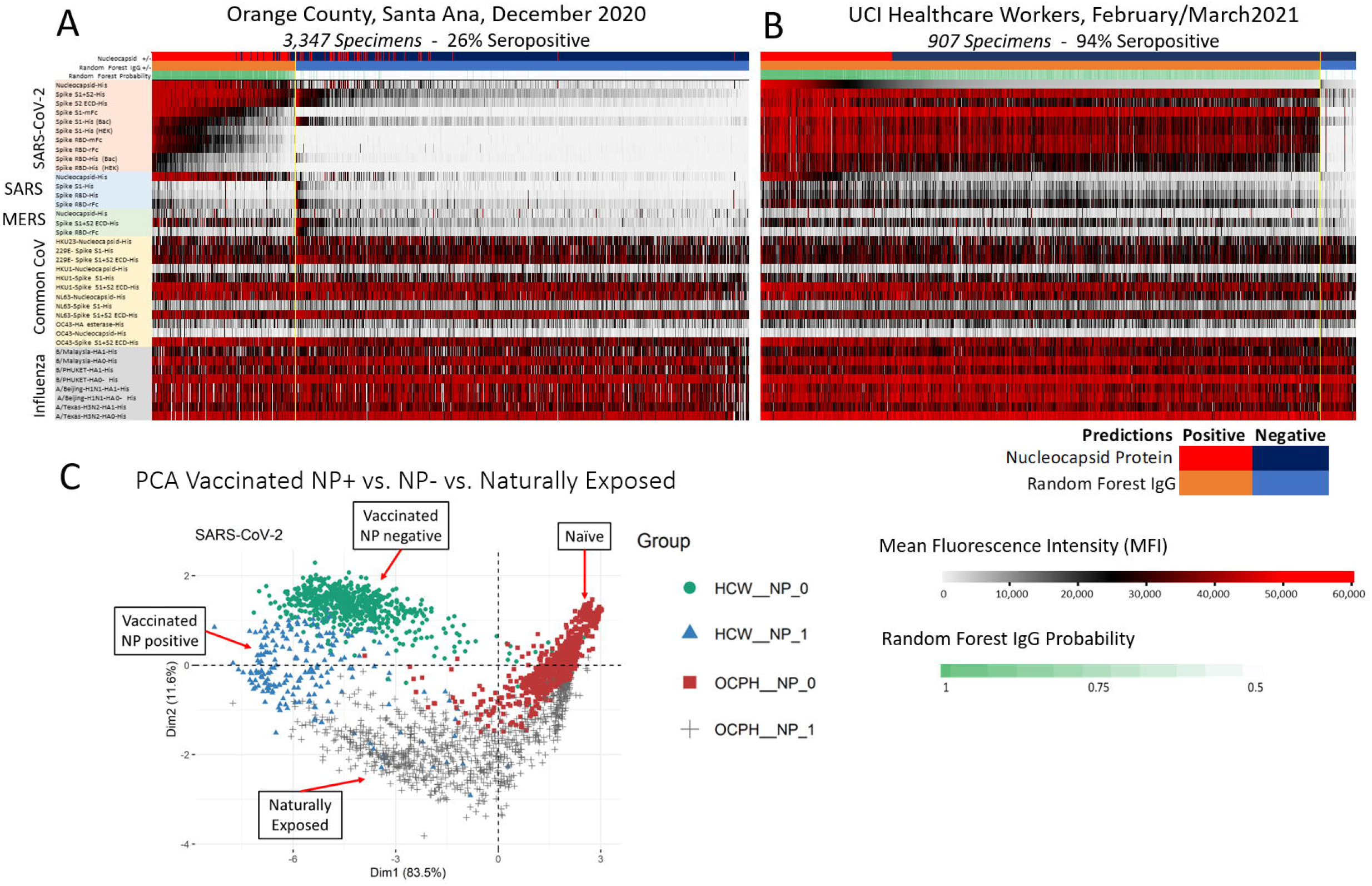
The heat maps show all of the IgG reactivity data from 3,347 pre-vaccination specimens collected from Santa Ana in December 2020 (**A**), and 907 post-vaccination specimens collected from the UCIMC in February (**B**). The 37 antigens are in rows and the specimens are in 3,347 columns for panel A and 907 columns for panel B. The level of antibody measured in each specimen against each antigen is recorded as Mean Fluorescence Intensity (MFI) according to the graduated scale from 0 to 60,000. Red is a high level, white a low level and black is in between. **Panel A**. Samples are classified as either SARS-CoV-2 seropositive clustered to the left (orange bar) or seronegative and clustered to the right (blue bar). Seropositive specimens recognize nucleoprotein and full-length spike. RBD segments are recognized less well. **Panel B**. Reactivity of specimens from 907 UCIMC HCW, 94% were vaccinated and seropositive. The heatmap shows that seropositive vaccinees in the HCW cohort can be classified into two groups, either seropositive for nucleoprotein or not, whereas the naturally exposed population (**panel A**) is uniformly seropositive for both nucleoprotein and full-length spike. (**C**) Principle component analysis of the protein microarray data in this study. The specimens fall into 4 distinct groups based on their reactivity against 10 SARS-CoV-2 antigens. Naturally exposed individual separate from unexposed naïves, the naturally exposed separate from the vaccinees, and the vaccinees separate into 2 groups depending on whether thy are seropositive for NP or not.

**Figure 4.**
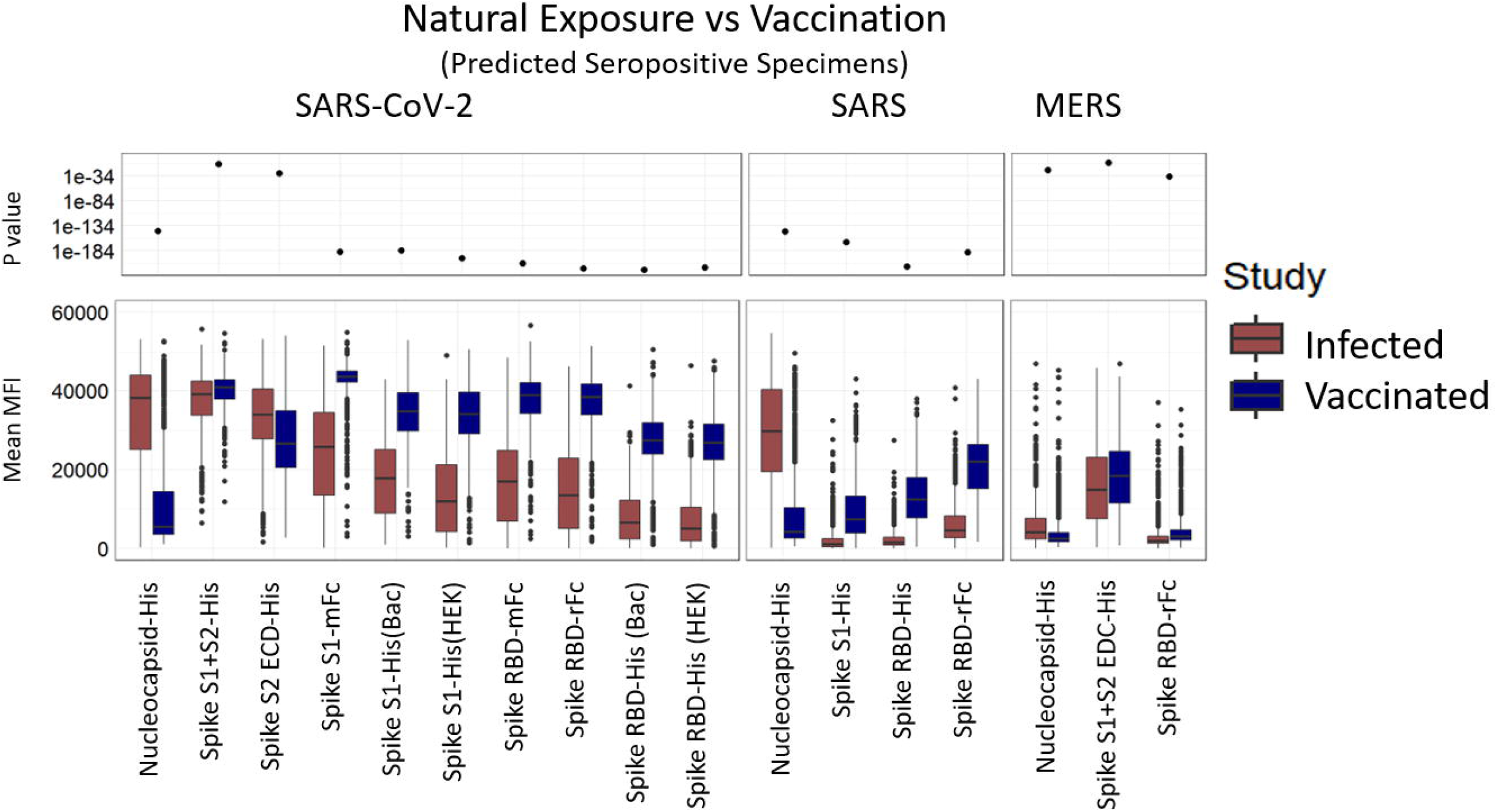
Mean MFI signals for each of the novel coronavirus antigens in the natural exposure cohort from Santa Ana in December 2020 (actOC) and the February/March 2021 vaccination group (HCW) are plotted. The figure shows that Ab responses against Spike RBD variants are significantly elevated in mRNA vaccinated people compared to naturally exposed individuals. Vaccination induces a broader and higher titer Ab response than natural exposure alone, so those who have recovered from COVID can be expected to benefit from the vaccination.

### Natural exposure and mRNA induced antibody profiles; anti-nucleocapsid Ab biomarker of natural exposure

Data from 3,347 specimens collected from Santa Ana residents in December 2020 are shown in the heatmap Figure 3A. The level of antibody measured in each specimen against each antigen is recorded as Mean Fluorescence Intensity (MFI) according to the graduated scale from 0 to 60,000. In order to assess the seroreactivity, we utilized a Random Forest based prediction algorithm that used data from a well characterized training set (pre-CoV seronegatives collected in 2019 and PCR-confirmed positive cases) to classify the samples as seroreactive or not seroreactive [6, 7]. This algorithm was constructed to classify SARS-CoV-2 serostatus using reactivity of 10 SARS-CoV-2 antigens to maximize sensitivity and specificity. With this machine learning algorithm, the samples were classified as either SARS-CoV-2 seropositive, grouped to the left, or seronegative and clustered to the right (Figure 3A). Seropositive specimens recognize nucleoprotein and full-length spike. RBD segments are recognized less well.

The heatmap in Figure 3B shows reactivity of specimens from 907 UCIMC healthcare workers collected in February and March after the vaccination campaign.; 93.8% were seropositive, of whom most were vaccinated. The anti-SARS-CoV-2 Ab reactivity induced by vaccination (Figure 3B) differs from the Ab profile induced by natural exposure (Figure 3A). The vaccine induces higher Ab levels against the RBD containing segments compared to the level induced by natural exposure in the Santa Ana cohort.

Since all adults in these cohorts are exposed to seasonal colds, influenza virus infections, and influenza vaccinations, all the individuals have baseline Ab levels against common-cold CoV and influenza. Thus, background Ab levels against all common CoV and influenza antigens are elevated in both the Santa Ana and HCW groups irrespective of whether they are COVID seropositive or not.

Principal component analysis using the reactivity to the SARS-CoV-2 antigens (Figure 3C) shows that seroreactive samples from the two study groups fall into two clusters (mainly along the 1^st^ dimension axis) indicating that the antibody response to the vaccine differs from the antibody response induced by natural infection. In addition, the heatmap (Figure 3B) clusters seropositive vaccinees into two groups based on whether they are seropositive for SARS-CoV-2 NP or not. The naturally exposed population (Figure 3) shows high reactivity to both SARS-CoV-2 NP and full-length spike (S1+S2). This is also evident in the PCA analysis which shows distinct clustering according to the reactivity to the nucleocapsid protein (NP, mainly along the Dimension 2 axis).

### mRNA vaccines induce higher Ab levels and greater Ab breadth than natural exposure to infection

Mean MFI signals for each of the novel coronavirus antigens in the Santa Ana natural exposure and the UCIMC vaccination healthcare workers groups are plotted in Figure 4. Natural exposure in seropositive people induces high antibody levels against NP, full-length spike (S1+S2) and the S2 domain. Antibodies against S1 and the RBD domains are lower. Vaccinated individuals have high Ab levels against full-length spike and the S2 domain of SARS-CoV-2 spike, and significantly higher antibody levels against S1 and the RBD domains compared to naturally exposed individuals. In natural exposure there was no significant cross-reactivity against SARS S1 or the RBD domains. Surprisingly, the vaccine induced significant cross-reactive Abs against the SARS spike and SARS RBD. Cross-reactivity against SARS NP and full-length MERS S protein is evident in both the natural exposure and vaccinated groups. These results indicate that antibody responses against spike RBD variants are significantly elevated in vaccinated compared to naturally exposed individuals. Vaccination induces a more robust antibody response than natural exposure alone, suggesting that those who have recovered from COVID benefit from the vaccination with stronger and broader antibody response.

### mRNA vaccines induce Abs that cross-react against SARS spike

Cross-reactivity of the SARS-CoV-2 NP antibodies induced by exposure to the virus, against NP from SARS is evident from the scatterplot in Figure 5A. The antibodies induced by SARS-CoV-2 infection react equally against NP from both SARS-CoV-2 and SARS. Cross-reactivity against SARS NP and full-length MERS S protein is also evident in both the natural exposure and vaccinated groups. However, significant cross-reactivity to SARS S1 and SARS RBD domains was only observed in the mRNA vaccine group.

**Figure 5.**
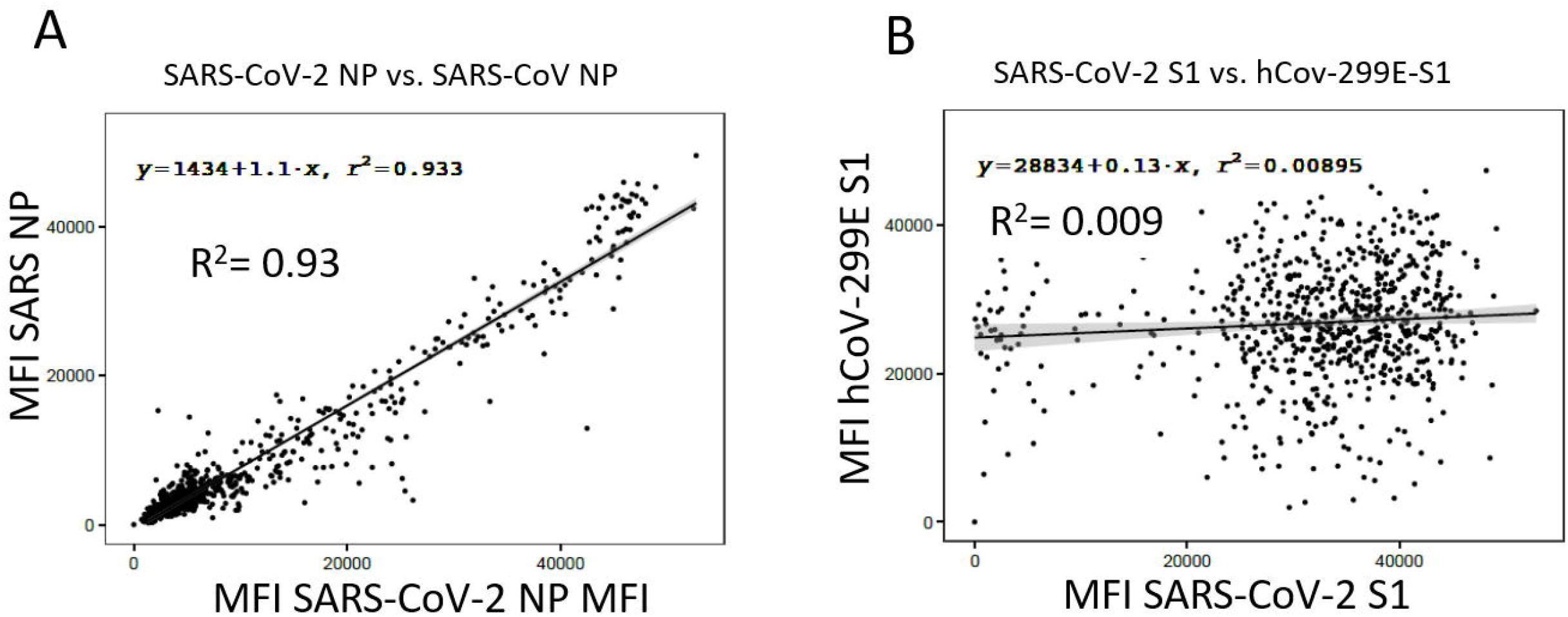
Scatterplots can be used to compare Ab reactivities of any 2 antigens on the COVAM array. **(A)** There are 920 seropositive specimens from Orange County residents. Ab reactivity against SARS-CoV-2 and SARS NP in this population are well correlated (R^2^= 0.93). Antibodies against NP from SARS-CoV-2 cross reactive against the NP from SARS. **(B)** Ab reactivities between SARS-CoV-2 S1 and hCoV-299E S1 are not correlated (R^2^= 0.009) so antibodies against SARS-CoV-2 S1 do not cross-react against S1 from hCoV-299E. The R^2^ value can be used as a metric to determine cross-reactivity between any 2 antigens.

This cross-reactivity can be shown using the reactivity correlation between the SARS-CoV-2 spike antigens and Non-SARS-CoV-2 antigens as a surrogate. As a representation, the correlation between two cross reactive antigens (SARS-CoV-2 nucleoprotein and SARS nucleoprotein) as well as two non-cross-reactive antigens (SARS-CoV2-S1 and hCoV-229E-S1) are shown in figure 5. The scatterplot returns an R^2^ value equal 0.93 indicating that NP antibodies induces by SARS-CoV-2 infection cross-react with SARS NP. Similarly, the Ab reactivity of SARS-CoV-2 S1 can be plotted against the common CoV 299E S1 producing an R^2^ value of 0.009 showing that they are not correlated and there is no significant cross-reactivity between these two S1 antigens. (Figure 5B).

There are 37 antigens on the COVAM and 702 pairwise comparisons. The R^2^ values for all pairwise comparisons are plotted on the correlation matrices in Figure 6. Figure 6A plots cross-reactivity of antibodies induced by natural exposure, and Figure 6B the cross-reactivity of antibodies induced by vaccination. Natural exposure induces SARS-CoV-2 NP antibodies that cross react with SARS NP. Anti-full length spike antibodies that cross-react with S2, but not against S1 and the RBD domains (Figure 6A, Green box). All of the anti-S1 Abs cross-react with the RBD domains. There is no cross reactivity evident against SARS S1 or SARS RBD (Figure 6A, Blue box). mRNA vaccination (Figure 6B) shares cross-reactivity of natural exposure. The mRNA vaccine also induces antibody against full length spike that cross-reacts with SARS-CoV-2 S1 and the RBDs (Figure 6B, Green box). In addition, the vaccine induced antibody against spike cross reacts with SARS S1 and RBD. (A complete list with all correlation coefficients can be found in the supplementary table X)

**Figure 6.**
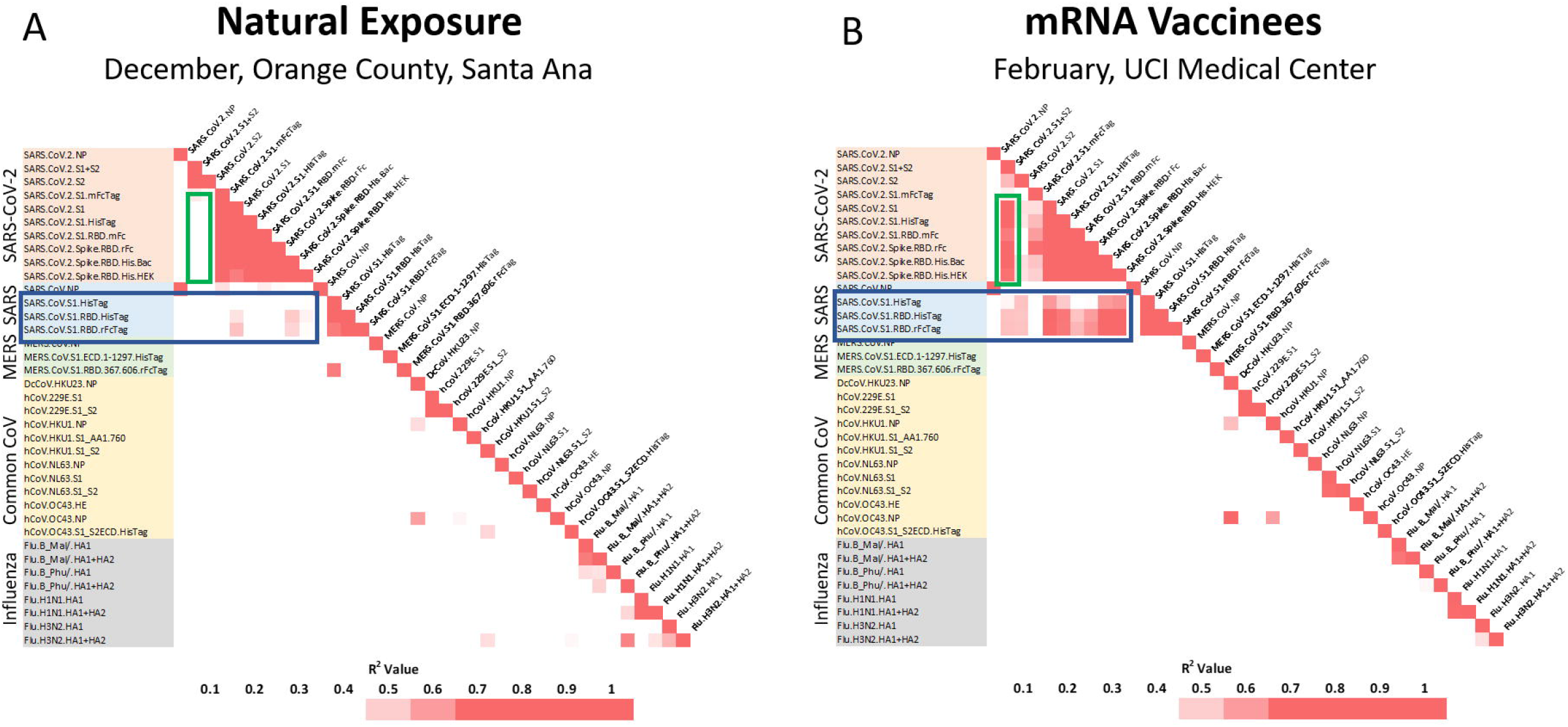
Correlation matrices with all pairwise comparisons between all antigens on the COVAM array were generated. The heatmaps represent a color scale of the r-squared of each pairwise comparison. On (**A**) is shown the correlation matrix for the Orange County group (actOC Natural exposure) and in (**B**) is shown the UCIMC vaccinated group. The mRNA vaccine induces cross reactive antibodies against SARS S1 and the RBDs (Figure B, Blue Box) and naturel exposure does not (Figure A) Similarly, vaccine induced antibodies against full length spike cross-react with SARS-CoV-2 RBD (Figure B, Green Box) and the natural exposure does not (Figure A).

As shown here and previous work from our group [6, 7] the specific antibody background reactivity to the novel coronavirus (SARS, MERS, and the SARS-CoV-2) is low in naïve populations and rises in response to the infection. However, during natural exposure, cross-reactivity was only observed between SARS-CoV-2 and SARS nucleocapsid proteins or MERS full length spike and SARS-CoV-2 S2 (or full length) was observed. Although it is possible to discover SARS-CoV-2 peptide epitopes that cross-react with peptide epitopes from common CoV [11], the results in Figure 6 emphasize the low level of cross reactivity against common CoV and flu conformational epitopes represented on the COVAM.

### Nucleocapsid protein is a biomarker associated with natural exposure

Unlike the natural exposure group that reacts uniformly to both nucleoprotein and full-length spike, vaccinees can be separated into two distinct groups of those who react to NP and those who do not. Natural exposure induces a dominant Ab response against the nucleocapsid protein (NP), but since NP is not in the vaccine, there is no vaccine induced response against it. In this way vaccinated people who had a prior natural exposure can be classified because they have Abs to NP. Vaccinated people who were never previously exposed lack Abs against NP and vaccinated healthcare workers can be separated into NP negative and NP positive groups.

The results in Figure 7 compare the Ab responses against the novel coronavirus antigens between the NP positive and NP negative vaccinees. Overall, it was observed that NP reactive individuals show a higher reactivity to the spike antigens, including cross-reactive from SARS spike, and a lesser degree MERS. This observation further supports the advice that people who were previously exposed will benefit from getting vaccinated as the antibody response can be further boosted by the vaccine.

**Figure 7.**
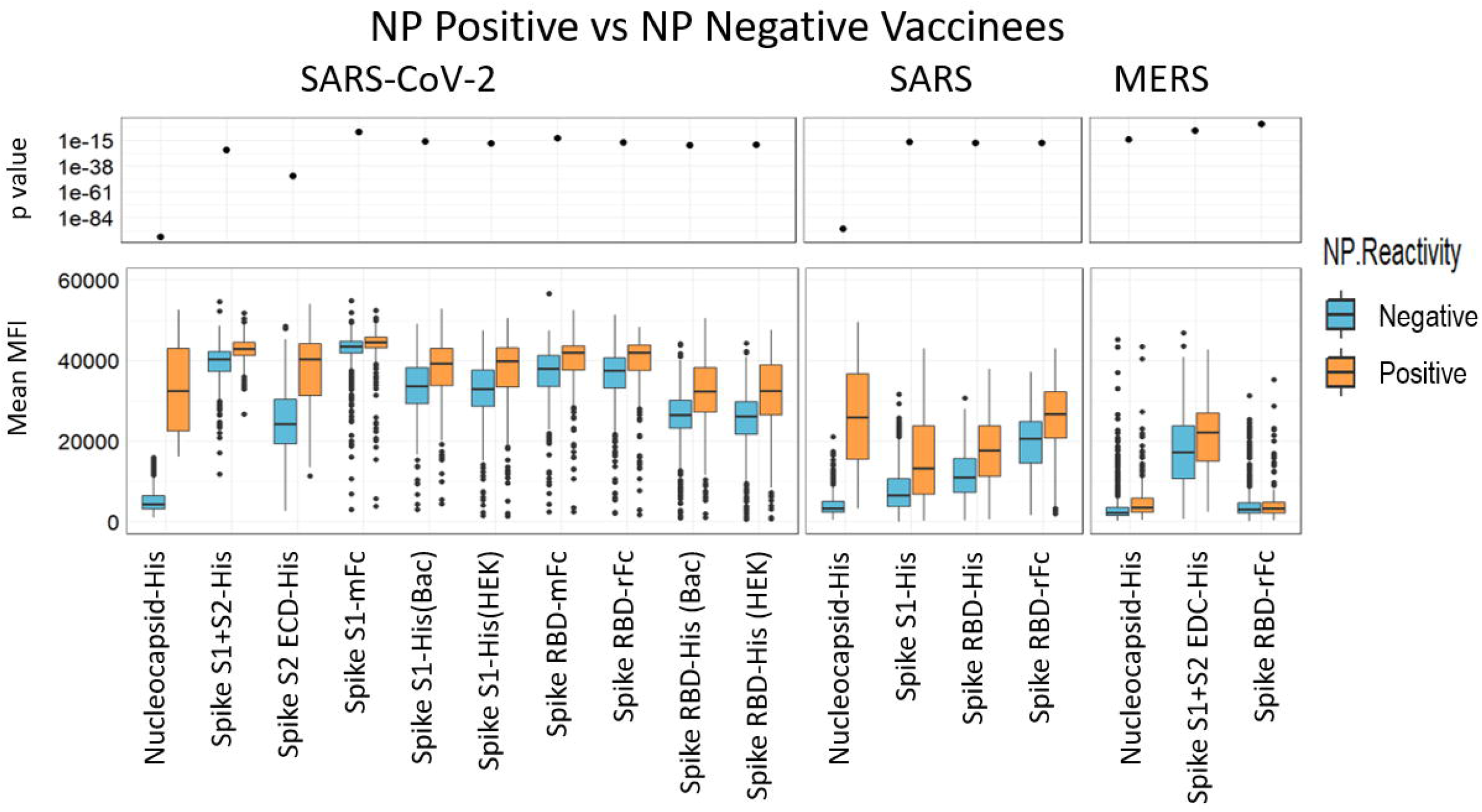
Unlike the natural exposure group that reacts uniformly to both nucleoprotein and full-length spike, vaccinees can be separated into two distinct groups, those who react to NP and those who do not. Natural exposure induces a dominant Ab response against the nucleocapsid protein (NP), but since NP is not in the vaccine, there is no vaccine induced response against it. In this way vaccinated people who had a prior natural exposure can be classified because they have Abs to NP. Vaccinated people who were never previously exposed lack Abs against NP. This data further supports the directive that people who are previously exposed will benefit by getting a boost against RBD.

### Progression of the prime and boost responses differ between individuals

Figure 8 shows results of longitudinal specimens taken at varying intervals from 9 individuals pre- and post-mRNA vaccination. Everyone received two doses of the vaccine, a prime and a boost roughly 4 weeks after the primary dose. The data show that the time course of development of the antibody response varies between each individual. There was no significant vaccine induced increase in NP reactivity as expected. The subjects showed either a plateau in the reactivity 5 to 10 days after the boost dose or a small decrease in reactivity. It is not yet clear whether this decrease is a sign of the waning antibody response.

**Figure 8.**
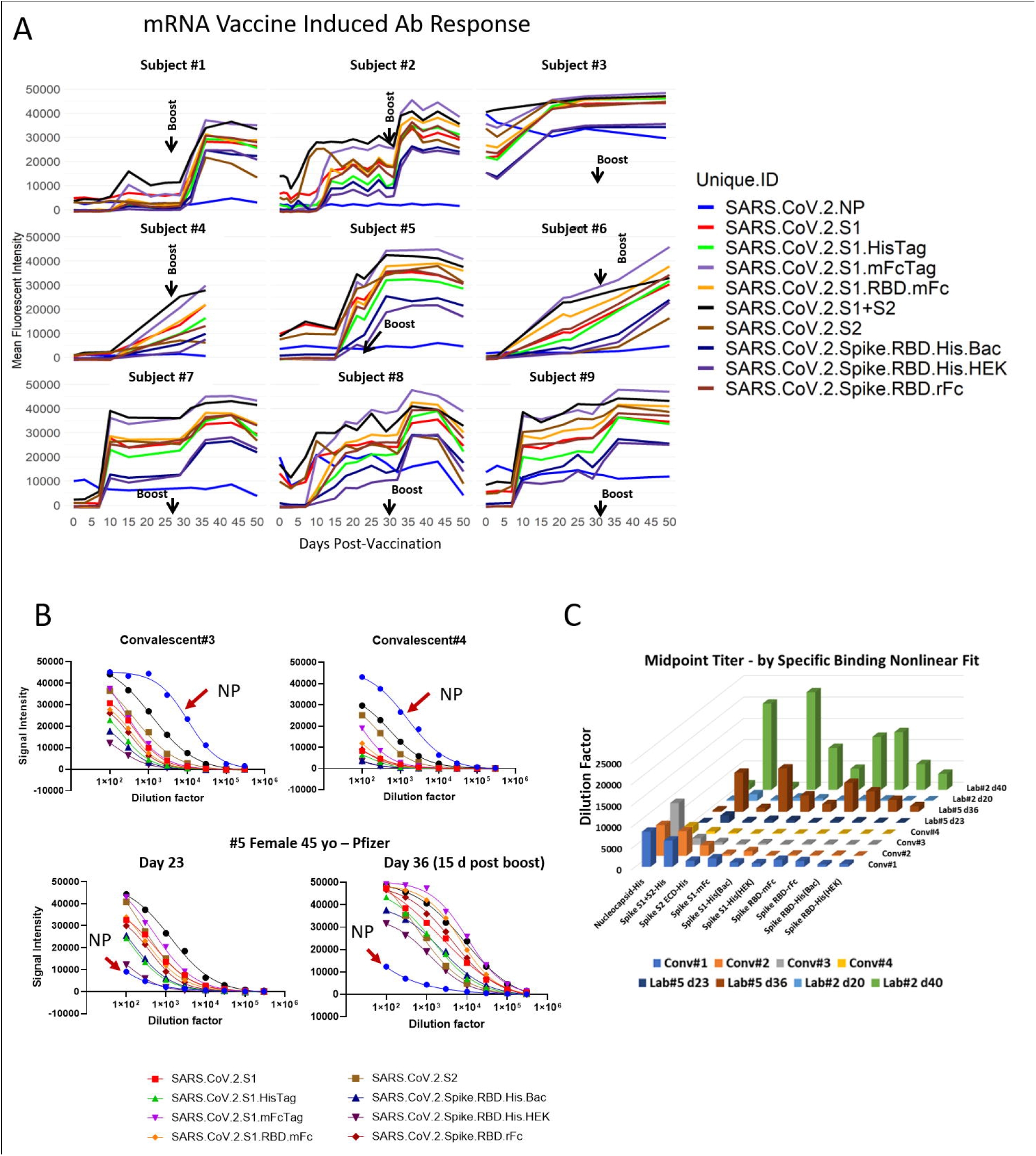
(**A**)Longitudinal specimens taken at weekly intervals from 9 individuals pre- and post-mRNA vaccination. Individuals differ substantially in their response to the prime. Five individuals had low baseline NP reactivity that did not change post-vaccination. Four individuals had elevated NP reactivity at baseline which also did not change significantly post-vaccination; subject #3 was a recovered confirmed COVID case. In this small group, higher baseline NP predicts a higher response after the prime. These results support a directive to get the boost in order to achieve more uniform protection within a population of individuals. (**B**)Convalescent plasmas from 2 recovered COVID cases, and pre- and post-boost specimens from Subject #5 were titered and the titration curves are shown. The curves are generated by making 8 half log serial dilutions of the plasmas before probing 8 separate COVAM arrays. These curves highlight the observation that high titers against NP are present in convalescent plasma that are lacking in the vaccinees. (Red Arrow). (**C**) The midpoint titers of 10 SARS-CoV-2 antigens from 4 convalescent plasmas and plasmas from 2 vaccinees after the prime and after the boost are plotted Convalescent plasmas vary in their titers against NP and full-length spike. The vaccinees lack Ab against NP and have significantly higher titers after the boost against all of the spike antigens compared to convalescent plasma.

Five individuals had low baseline NP reactivity that did not change post-vaccination. Four individuals had elevated NP reactivity at baseline which did not change significantly post-vaccination, and one of these individuals was a confirmed recovered COVID case. Subject #1 had a weak response to the prime and a stronger response to the boost. #2 responded with a strong reactivity to both the prime and the boost with a clear increase in antibody levels for the spike variants. #3 is a recovered confirmed COVID-19 case. As expected, this individual showed an elevated baseline Ab reactivity against NP and all of the SARS-CoV-2 variants. After the first dose, the individual showed an increase in antibody reactivity, however, no further increase was observed after the boost dose. #4 responded slowly to the prime. Subjects #7, #8 and #9 had elevated NP at baseline and responded rapidly to the prime without significant further increase after the boost.

### Anti-spike Ab titers induced by the mRNA vaccine are higher than those induced by natural exposure

COVAM measurements taken at a single dilution of plasma can be used as a parameter to compare relative antibody titers between individual specimens. This is useful for high throughput studies and allows for the probing of thousands of samples in a relatively short time, with minimum staff, and can provide fast and inexpensive data for epidemiology studies to quantify virus exposure levels. However, to obtain a more precise measure of antibody levels, samples can also be titered by serial dilution. In Figure 8B, 2 convalescent plasmas from recovered COVID cases, and pre- and post-boost vaccination plasmas from Subject #5 were titered. The curves are generated by making 8 half-log serial dilutions of the plasmas before probing the COVAM arrays. These curves highlight the observation that high titers against NP are present in convalescent plasma that are lacking in the vaccinees.

Figure 8C plots the midpoint titers of 10 SARS-CoV-2 antigens in 4 convalescent plasmas and pre- and post-boost plasmas from 2 vaccinees. As expected, convalescent plasmas vary in their titers against both NP and full-length spike. The convalescent plasmas #1 and #2 showed a higher midpoint titer for both NP and full length spike when compared to the plasmas #3 and 4. Both vaccinees showed no Ab reactivity against NP before and after immunization. Although both individuals showed low antibody titer against SARS-CoV-2 antigens right after the primary immunization, both showed significantly higher titers after the boost against all of the spike antigens including S1 and the RBDs, compared to convalescent plasma (Figure 8C). A summary of the midpoint titers is available in supplementary Table 1.

## Discussion

In this study, we compared antibody responses induced afterSARS-CoV-2 natural exposure with the responses induced by the mRNA vaccines. Pre-vaccine natural exposure data was obtained from two large serial cross-sectional surveys of residents from Orange County and the city of Santa Ana, CA, [10] and from mRNA vaccinated healthcare workers at the UCI Medical Center participating in an aggressive vaccination campaign. Within weeks of administration, the mRNA vaccines induced higher Ab levels against spike proteins than observed after natural exposure. These results coincide with equally remarkable clinical trial data showing rapid induction of mRNA protective efficacy on a similar timescale. [1, 2]

The UCI Medical Center achieved a very rapid introduction of the vaccine beginning on December 16, 2020. Within 5 weeks 78% of the individuals tested were seropositive for spike and 3 months later 99% of a March 2021 cross sectional sample was positive. These results illustrate the high vaccine uptake and the extent of antibody response to the vaccine in this population.

mRNA vaccines induce higher Ab levels and greater Ab breadth than natural exposure to infection and differences were particularly notable against the RBD domain. Out of a collection of 3,473 specimens collected from the Santa Ana Cares study in December 2020 we classified 920 as seropositive due to natural exposure before the vaccine was introduced. In February we had a similar number of vaccine induced seropositive healthcare workers. The virus uses the spike RBD domain that binds to the ACE2 receptor on respiratory cells to enter and infect them. Vaccinated individuals had significantly elevated Ab levels against RBD domain segments, supporting the protective immunity induced by this vaccine as previously published. [1, 2] To account for this difference between natural exposure and the vaccine, the virus may have evolved to conceal the RBD epitope to evade immune recognition. The mRNA vaccine produces a protein conformation that better exposes the RBD epitope to the immune system.

In addition to inducing increased Ab levels against SARS-CoV-2 RBD, the mRNA vaccine induced cross-reactive responses against SARS spike and SARS RBD. Conversely, natural exposure did not induce a cross-reactive response against the SARS spike and SARS RBD. This result can be interpreted based on immune selection pressure. The weak anti-RBD response induced by natural exposure may provide a mechanism for new variants to enter the population. Importantly, the mRNA vaccine induces a marked cross-reactive response against SARS spike, indicating that the mRNA vaccine adopts a conformation that presents cross-reactive epitopes to the immune system. This effect of the mRNA vaccine to induce cross-reactivity against diverse CoV strains is encouraging, providing further evidence that it may be effective against emerging virus variants.

Antibodies induced by natural exposure against the NP from both SARS-CoV-2 and SARS is concordant with an R^2^ value of 0.85. This may indicate a relative lack of selective pressure on this antigen during evolution of these two CoV species. Conversely, the anti-spike response induced by natural exposure does not cross-react against SARS spike or SARS RBD domain indicating immune selection pressure across these strains because of the importance of this epitope in the infection process.

Anti-nucleocapsid Ab is a biomarker of natural exposure to SARS-CoV-2 and can be used to distinguish individuals in a vaccinated population who have been previously exposed to the virus. The nucleoprotein is not present in currently used vaccines. Our data also suggests that people who have had a prior exposure to the virus mount a stronger immune response to the vaccine than those whose immune response has not yet been primed by a previous exposure or vaccination.

These results may also have relevance for both the dose response hypothesis and regarding herd immunity. Several authors have suggested that disease outcomes may be related to the dose inoculum, with individuals being exposed to inocula with higher virus loads potentially having more severe disease outcomes. [12] While the currently used vaccines in this setting do not rely on viral materials, they do offer a glimpse into controlled high-level exposure to proteins that are specific to SARS-CoV-2. Our results show that individuals who have been vaccinated mount higher across-the-board antibody responses than those who have been exposed to variable viral inocula (i.e. through natural exposure). Second, the variable antibody responses among the pre-vaccine population may also indicate that immune responses to natural infections are not as strong as those among individuals who have been vaccinated. This could also indicate that immunity from naturally acquired infections is not as strong as that acquired from vaccination, with potential relevance for reaching and maintaining herd immunity. We should not assume that previously infected individuals are immune or that they cannot transmit the virus.

The original influenza nucleic acid vaccination report published nearly 30 years ago, used the nucleoprotein antigen from influenza because it was conserved across influenza subtypes and it would therefore be a more universal vaccine [13]. The experiment was successful, it was universally effective across diverse strains, and it implicated a cell mediated component, killing of infected cells, in the observed efficacy. As reported for influenza, a more universal SARS CoV vaccine may include the nucleocapsid protein antigen.

Individuals differ in the progression of response to the mRNA prime and boost. Some have a weak response to the prime and experience a substantial effect of the boost. To account for these differences, the group of vaccinees that are NP positive also have significantly higher vaccine induced responses than the NP negative individuals. This effect is also evident from the small sample of longitudinal specimens we collected from lab members, those with elevated baseline NP reacted more rapidly against the antigens. In the small sample of logitudinal specimens, anti-spike Ab titers induced by the mRNA vaccine are higher than those induced by natural exposure

Serological assays for SARS-CoV-2 are of critical importance to identify highly reactive human donors for convalescent plasma therapy, to investigate correlates of protection, and to measure vaccine efficacy and durability. Here we describe results using a multiplex solid phase immunofluorescent assay for quantification of human antibodies against 37 antigens from SARS-CoV-2, other novel and common coronaviruses, and influenza viruses that are causes of respiratory infections. This assay uses a small volume of blood derived from a finger stick, does not require the handling of infectious virus, quantifies the level of different antibody types in serum and plasma and is amenable to scaling. Finger stick blood collection enables large scale epidemiology studies to define the risk of exposure to SARS-CoV-2 in different settings. Since the assay requires 1 microliter of blood it is also practical for monitoring immunogenicity in small animal models. After probing more than 8,000 pre- and post-vaccination specimens our results confirm that the mRNA vaccine can be used in an aggressive and targeted vaccination campaign to immunize large groups within a matter of weeks.

There are stark differences between actionable interpretation of molecular PCR results and the serological results like those reported here. PCR tests answer the question whether a person has virus in their respiratory secretions as a confirmatory test accounting for the cause of COVID symptoms. It is a useful test in settings where there is high incidence of active infection, patients experiencing symptoms, household contacts, and for contact tracing. Serological tests address different questions of whether the individual has an immune response to the virus, could I have immunity to the COVID 19 virus, how long does it last, do I need the vaccine if I had COVID, can I go to work yet, which vaccine is better, and when do I need another shot.

The concept of nucleic acid vaccines appeared 30 years ago after it was shown that plasmid DNA and RNA could be injected into mouse skeletal muscle tissue in vivo and the encoded transgenes were expressed at the injection site. [14, 15] After intramuscular (IM) injection of a plasmid encoding HIV gp120, induction of anti-gp120 Abs was reported[16]. That was followed by a 1993 report showing efficacy of an influenza nucleic acid vaccine in a rodent model[13]. This was a nucleocapsid based nucleic acid vaccine that induced cross-subtype protection against both group 1 and group 2 viruses (A/PR/8/34 (H1N1) and A/HK/68 (H3N2)). The utility of cationic lipids for gene delivery was discovered and reported in 1987 [17] and synthetic self-assembling lipoplexes for gene delivery described[18–20]. These results spawned a branch of gene therapy science, and an NIH study section, Genes and Drug Delivery (GDD) was established in 2002 that continues to support this research emphasis. Since then synthetic gene delivery system research and nucleic acid vaccine science has flourished.

DNA vaccines were the first nucleic acid vaccines to be manufactured and tested on a pharmaceutical scale [21, 22]. The mRNA vaccines that are being distributed so widely today may seem to have suddenly emerged, but there has been 30 years of scientific discovery, discourse and development, work from hundreds of scientists, numerous biotechnology companies and billions of public and private dollars invested enabling this effective response with a vaccine at this moment.

## Methods

### COVID seroprevalence surveys in Orange County, California

Here we analyzed data from ongoing serologic surveys of healthcare workers (HCW) from the University of California Irvine Medical Center (UCIMC, Orange County, CA, USA) and from residents of the Orange County community. The first community survey (actOC) conducted in July of 2020, was county-wide, and recruitment was done via a proprietary phone list. This survey of 2,979 individuals was meant to be representative of the age, ethnicity, and socio-economic makeup of the county (detailed in [10]). The results of this county-wide survey indicated that the city of Santa Ana was a COVID-19 hotspot, especially on the Hispanic population. Surveillance of reported cases and test positivity corroborated this finding. A second, seroprevalence survey was then conducted in Santa Ana as the Santa Ana Cares study in December of 2020. Recruitment of 3347 individuals for this second survey was done using randomized house sampling within cenus tracts coupled with a community engaged campaign with support from Latino Health Access (a community-based health organization that has been based in Santa Ana for over 2 decades, https://www.latinohealthaccess.org/). Analysis of the second seroprevalence survey is ongoing. While the first survey was county-wide, the serological test positivity reported in this analysis come from zip codes in Santa Ana alone.

Samples were also collected from the UCIMC longitudinal HCW study in May and December 2020. An aggressive and comprehensive mRNA vaccination campaign started at UCIMC on December 16 2020 and 6,724 HCW were vaccinated in 3 weeks. Three additional cross-sectional samples were taken at end of January, February, and March 2021.

A Coronavirus Antigen Microarray (COVAM) was used to measure antibody levels against 37 antigens from coronaviruses and influenza. COVAM measurements taken at a single dilution of plasma can be used as a parameter to compare relative Ab titers between individual specimens against each of the individual 37 antigens. The COVAM contained 10 SARS-CoV-2, 4 SARS, 3 MERS, 12 Common CoV and 8 influenza antigens. (Figure 1) Samples were probed and analyzed on the COVAM and each individual was provided with the results of their test (Supplementary Section) according to the IRB protocol. [6].

## Supplementary Methods

### Coronavirus Antigen Microarray (CoVAM) Report

This document describes the pipeline used to analyze the COVAM array and generate the individual reports.

### Step 1: Data pre-processing

The first step of the analysis is importing all data into the R environment. The sample set containing the known negative and known positive controls, here named “Control Set”, is loaded separately from the sample set being analyses.

Following this step, to prevent errors when addressing specific columns, or samples, all spaces are removed both from the column names from all data sets imported, as well as from the Unique sample IDs reference from the meta data files.

On the data processing steps, the following are performed:

From the raw data, the signal to noise ratio (SNR) is calculated. The SNR is calculated as the median signal intensity of a given spot divided by the background signal of the vicinity surrounding area. For the quality check purposes, the mean SNR is Calculated only for spots with MFI over 20,000. Samples with a mean SNR below 2 are flagged for further visual inspection or for reprobing.

After calculating the men SNR, the control spots are then assessed. First, for each sample, and each antigen (printed in triplicates), the first and third quartile as well as interquartile range (IQR) are calculated for the control spots. An upper MFI limit of 1.5 times the IQR over the third quartile and a lower limit of 1.5 times the IQR bellow the first quartile are defined. Spots outside this range are removed and replaced with the mean MFI of the remaining replicates of the spot.

Next, a similar approach is applied to flag samples for which the overall control spots distribution is out of range (2*IQR + third Quartile for the upper limit and first quartile – 2*IQR for the lower limit). For this, all controls spots of a given sample are used. Out of range samples are flagged for further visual inspection or reprobing.

Finally, the printing buffer background reactivity is subtracted from each spot and the samples are normalized.

### Step 2: Normalization

Data normalization is performed in two steps. First The control spots are normalized against the training set using the Quantile Normalization method. This allows to calculate a normalization factor that will be used to rescale the data to match the training set and preserving the individual reactivity diversity. After normalizing the control spots, their sum is calculated. A rescaling factor is calculated by dividing the sum of the normalized control spots of the training set by the sum of the normalized control spots of each sample. The resulting factor is then multiplied by the reactivity of each spot resulting in a rescaled data frame. The mean reactivity of the normalized data is then calculated.

### Step 3 a: Prediction models

Previous to the sample analysis, the prediction models were constructed using a sample set composed by samples with known diagnosis for COVID-19. These samples are both Negative controls (samples collected before the pandemic) and Positive controls (Samples from individuals diagnosed for COVID-19 by PCR). This control set is heer referred to as Training Set.

The Construction of the prediction models was performed as following.

1. Data is pre-processed and normalized as described above.
2. The reference data set was decomposed into a vector using the function ‘unmatrix’ from the package gData (version 2.18.0).
3. A mixture model is calculated for the vector using the function ‘normalmixEM’ from the package ‘mixtools’ (version 1.2.0).
4. A cutoff is then calculated as 3 standard deviations over the mean of the negative signal curve.
5. Wilcox test for each antigen was performed comparing the positive controls and negatives control, considering significant, antigens with p < 0.05.

following the selection of seropositive antigens, an optimal predictive combination of these antigens was selected. (that left us with 7 antigens as seropositive for IgG, and 8 for IgM).

The selection was performed as follows:

1. For every possible combination of the seropositive SARS-CoV-2 antigens from 1 all (7 for IgG and 8 for IgM), the reference set was randomly divided into a training and a testing sets at a 70%/30%ratio.
2. A logistic regression was generated using the reference set. The regression was generated using the function ‘glm’ of the ‘stats’ package (version 4.0.0).and a ROC curve was calculated (package pROC version 1.16.2).
3. The optimal coordinates of the ROC curve were obtained based on the ‘youden index’, by prioritizing the specificity.
4. The coordinates were obtainded using the function ‘coords’ from the pROC library. The coordinates are obtainded in a table format with each row containing a regression threshold and its related specificity and sensitivity.
5. The coordinates were then subset to represent specificities of 0.95 or higher. A threshold was then defined as the threshold on the coordinate with the highest specificity on the subset.
6. A logistic regression was then calculated using the testing set and each sample classified as negative or positive by comparison with the threshold.
7. A confusion matrix was calculated by comparing the predicted outcomes and the known classifications (“known negative” or “Known positive”) and the prediction specificity and sensitivity stored into a vector.
8. This analysis was repeated 1000 times and the sensitivity and sensitivity calculated as the mean predicted performance of all repetitions.

The performance outcome for each antigen combination was analyzed and a selection of the best performing combinations was made based on the specificity and sensitivity. The selected candidates were then tested using the full reference sample set. The test was performed as follows:

1. A logistic regression for each antigen combination candidate using the full reference set. Then a ROC curve was calculated and the coordinate table with all curve points was obtained.
2. The coordinates of each candidate were compared in order to select the candidate with the highest sensitivity, given a fixed specificity of 1 (100%).

In addition to the logistic regression model, a Random Forest model was constructed using all reactive antigens.

### Step 3 b: Reports

After Data Normalization, the predictions models, constructed as described above, are loaded and reactivity predictions are performed using Random Forest and Logistic Regression for the multi antigen combinations. In addition to the multi antigen predictions, a prediction for each single SARS-CoV-2 antigen was performed for every sample, for both IgG, and IgM. These predictions were performed using the threshold calculated using the optimal ‘youden’ index. Every sample can be classified as reactive or not reactive for each single SARS-CoV-2 antigen.

The report phase consists on the output of single pdf files with the individual subject predictions and interpretation. The file consists on a brief explanation of the array on the first page, as well as some information on the performance of the array with the current settings. In addition, on the first page there is a short disclaimer of the scope and limitations of the assay.

The second page consists of a table for all the SARS-CoV-2 antigens with their ROC predictions. These predictions are for a qualitative understanding of one’s reactivity and may not directly correlate with the multi antigen prediction.

The Multi antigen prediction, or the sample classification into the three reactive groups, is presented also on a short table displaying the prediction of IgG and IgM separately. The overall sero-reactivity of the sample to all antigens is depicted on two graphs on the second page. One showing the reactivity for IgG and one for IgM.

On each graph, the individual’s reactivity is represented as dots with its standard errors. For reference, a red line representing the positive control mean reactivity with its confidence interval, as well as a blue line representing the negative controls mean reactivity with its confidence interval are also plotted.

## Acknowledgements

This work was supported by the HDTRA1-16-C-0009, HDTRA1-18-1-0035, HDTRA1-18-1-0036 and a University of California, Irvine CRAFT-COVID grants. The findings and conclusions in this report are those of the authors and do not necessarily represent the official position or policy of the funding agencies and no official endorsements should be inferred.

## Author contributions

The coronavirus antigen microarray (COVAM) was designed by S. Khan and P. Felgner and was constructed by R. Nakajima, A. Jasinskas and R. Assis at UCI. The specimens probing on the COVAM was performed by A. Jain at UCI. Data analysis was performed by R. de Assis at UCI. The manuscript and figures were prepared by Philip Felgner and R. de Assis with input and approval from all other authors.

## Supplementary Figure Legends

**Supplementary Figure 1.** Mean MFI signals for the common coronaviruses and Influenza antigens in the natural exposure cohort from Santa Ana in December 2020 (actOC) and the February 2021 vaccination group (HCW) are plotted. The figure shows that Ab responses against common cold antigens are not significantly different in both populations. A realative higher reactivity was for the UCIMC group was observed for the influenza antigens.

**Supplementary Figure 2.** The general analysis pipeline consists of three main steps: the preprocessing, the normalization and then the statistical prediction analysis. The preprocessing includes steps like calculation the Signal to Noise Ratio (SNR) and determine if a sample needs to be further checked or re-assayed (due to the background reactivity levels). If successful, samples are successful analyzed for their SNR, the controls spots are checked to remove outlier spots that could skew normalization. Then, the distribution of the control spots is analyzed and low-quality samples (for which the control spots deviate from the expected) are flagged to be re-assayed. Then the samples are normalized, and the mean fluorescence intensity calculated from the average of the 3 replicates in the array. After normalization, a machine learning based algorithm is used to classify each sample as reactive or not reactive to SARS-CoV-2 (using multiple antigens) as well as to individual antigens. Then, individual reports are generated for each sample (this can be in the form of individual pdf files that may be delivered to the subject).

**Supplementary Figure 3.** After the machine learning classification of each sample individual pdf files containing the results can be generated. The panels in the figure are representative of a typical negative (or non-reactive) result (left panel) and of a typical positive (Reactive) sample (on the right). The data printed on the reports are basic reactivity classification for the SARS-CoV-2 antigens (Only reactive and Non-reactive denominations are given). As well as the machine learning classification (multi antigen classification) denominations. For the multi antigen classification, the results from the logistic regression as well as the results from random forest, as well as the random forest probabilities are given. The multi antigen classification is the main result and is the one used to classify an individual as exposed, or reactive to SARS-CoV-2 as individual antigens alone have a much lower performance in the classification.

Finally, since the COVAM is composed of multiple viruses, the reactivity to the entire array is given to both IgG and IgM. This reactivity is given as the normalized mean florescence intensities and as a reference, the confidence intervals of a known control set of samples (known positives red line and red bands and known negatives blue line and blue bands) are given. Although these reports give a much more comprehensive view of an individual’s reactivity status to SARS-CoV-2, they are intended mainly as a guidance as the COVAM array is not approved by the FDA as a diagnostic test.

