## Supplementary figures and images for "Distinct SARS-CoV-2 Antibody Responses Elicited by Natural Infection and mRNA Vaccination"

### Supplementary_Figure1

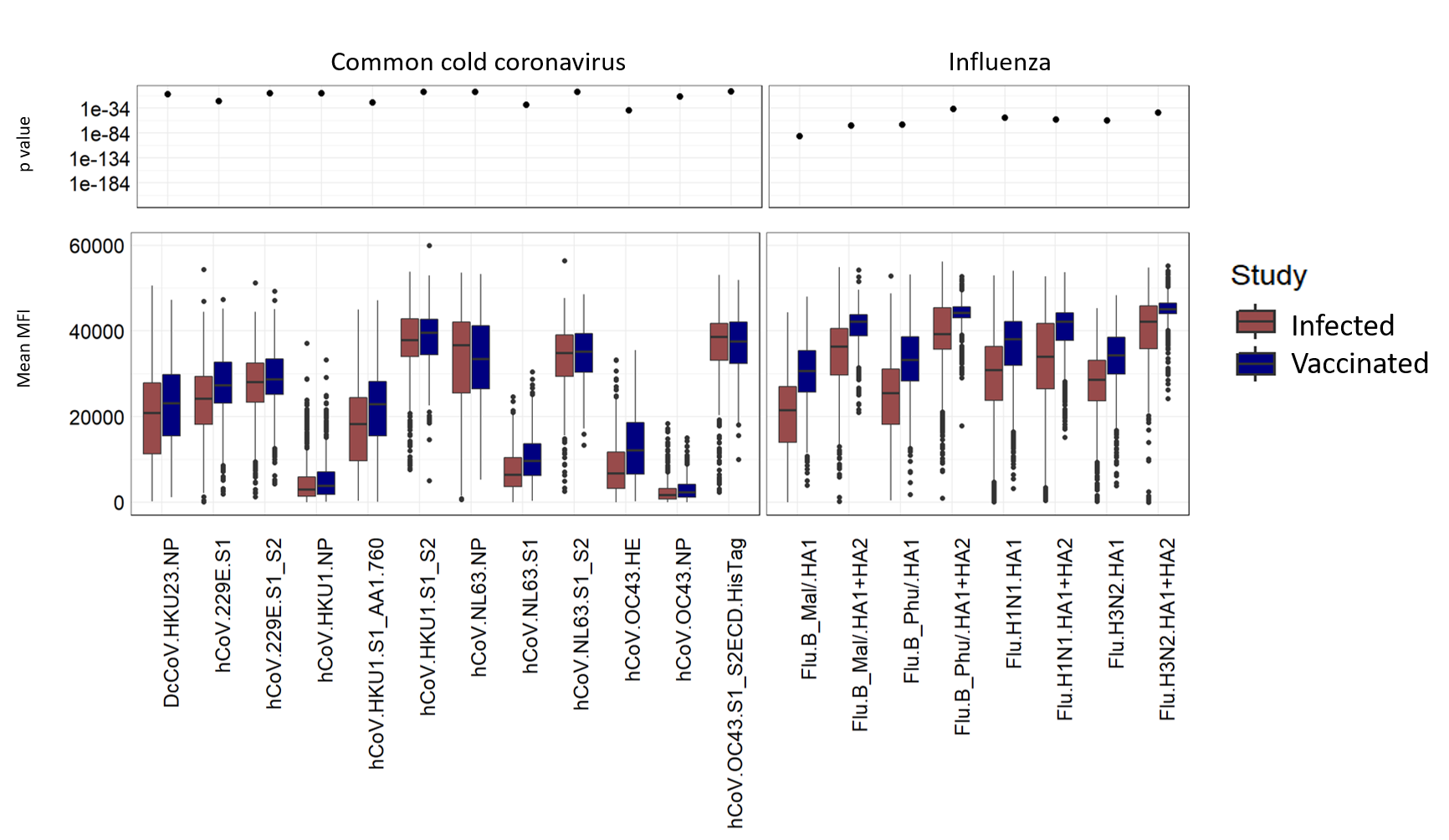

### Supplementary_Figure2

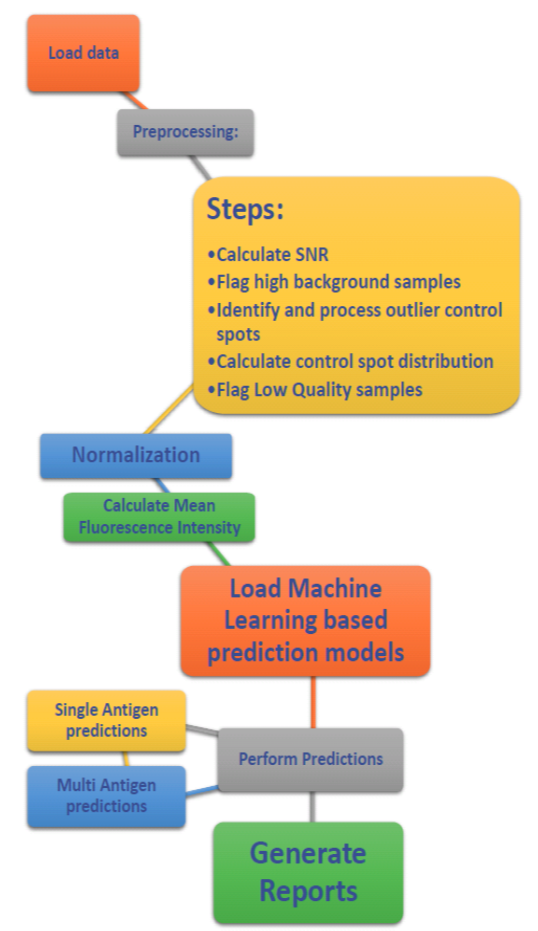

### Supplementary_Figure3

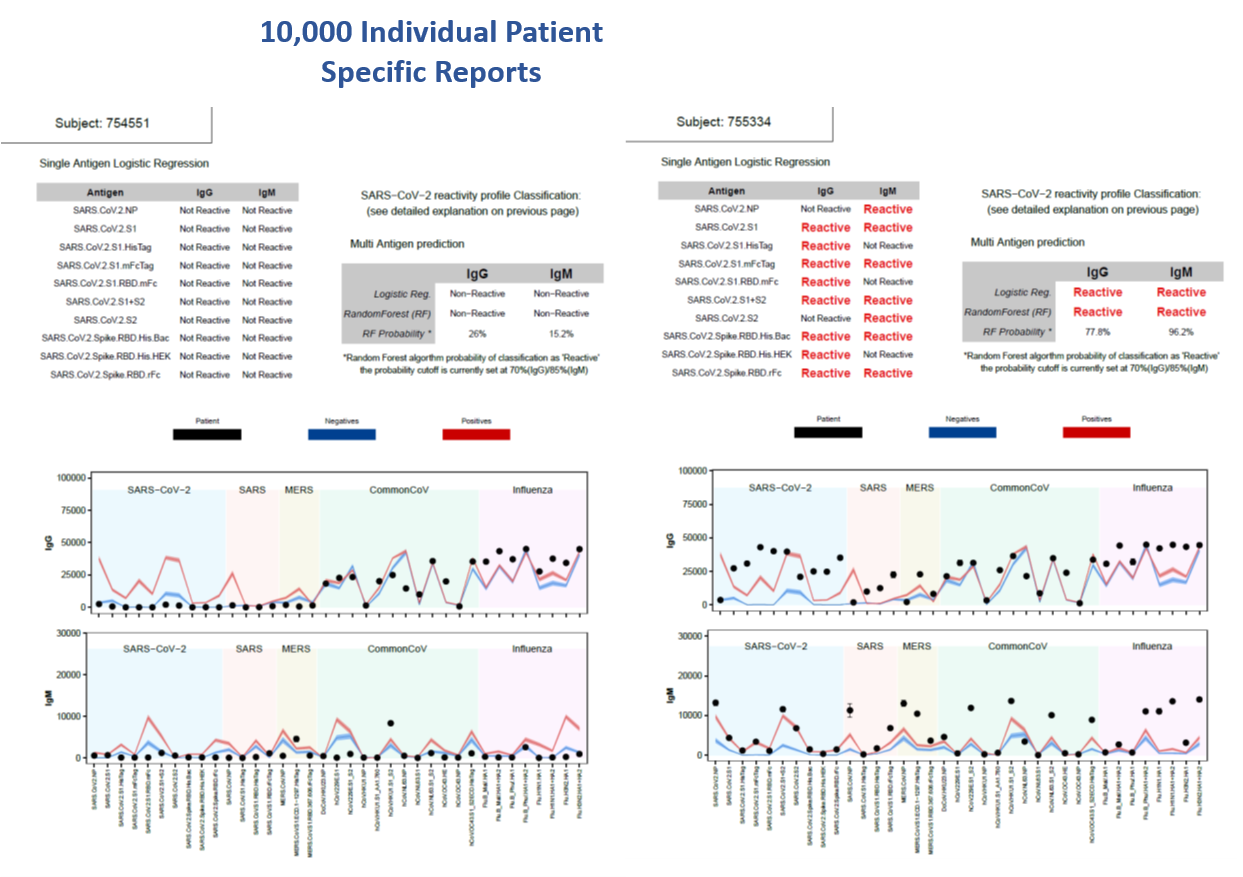

### Supplementary_Table1

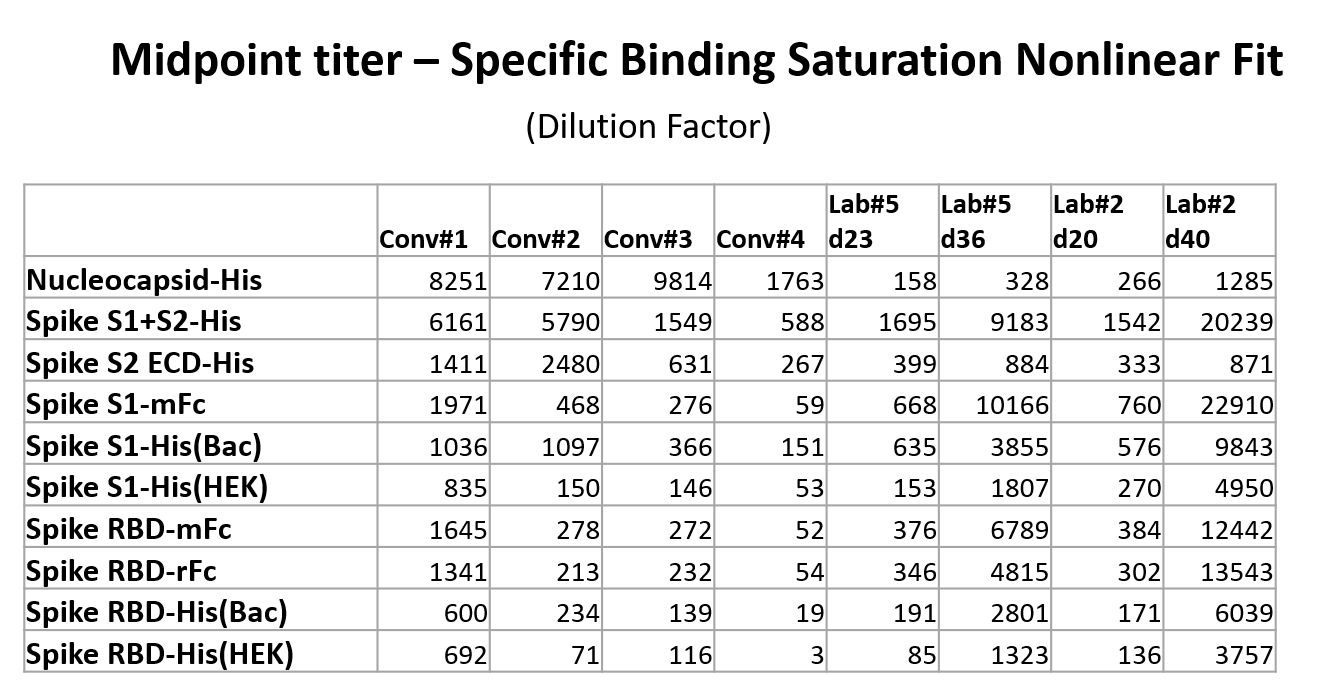
